# Customizable high-throughput platform for profiling cofactor recruitment to DNA to characterize cis-regulatory elements and screen non-coding single-nucleotide polymorphisms

**DOI:** 10.1101/2020.04.21.053710

**Authors:** David Bray, Heather Hook, Rose Zhao, Jessica L. Keenan, Ashley Penvose, Yemi Osayame, Nima Mohaghegh, Trevor Siggers

## Abstract

Determining how DNA variants affect the binding of regulatory complexes to cis-regulatory elements (CREs) and non-coding single-nucleotide polymorphisms (ncSNPs) is a challenge in genomics. To address this challenge, we have developed CASCADE (<u>C</u>omprehensive <u>AS</u>sessment of <u>C</u>omplex <u>A</u>ssembly at <u>D</u>NA <u>E</u>lements), which is a protein-binding microarray (PBM)-based approach that allows for the high-throughput profiling of cofactor (COF) recruitment to DNA sequence variants. The method also enables one to infer the identity of the transcription factor-cofactor (TF-COF) complexes involved in COF recruitment. We use CASCADE to characterize regulatory complexes binding to CREs and SNP quantitative trait loci (SNP-QTLs) in resting and stimulated human macrophages. By profiling the recruitment of the acetyltransferase p300 and MLL methyltransferase component RBBP5, we identify key regulators of the chemokine CXCL10, and by profiling a set of five functionally diverse COFs we identify a prevalence of ETS sites mediating COF recruitment at SNP-QTLs in macrophages. Our results demonstrate that CASCADE is a customizable, high-throughput platform to link DNA variants with the biophysical complexes that mediate functions such as chromatin modification or remodeling in a cell state-specific manner.

## Main

Determining the impact of genetic variation on cis-regulatory elements (CREs), such as enhancers and promoters that control gene expression, remains a challenge in modern genomics. Genome-wide association studies (GWAS) have identified thousands of single-nucleotide polymorphisms (SNPs) associated with human diseases, but the causal variants and their biological effects remain largely unknown^1–3^. Variants underlying disease risk often function by altering CRE function and gene expression. For example, >50% of causal SNPs for autoimmune diseases are non-coding SNPs (ncSNPs) mapping to immune gene enhancers^4^. Therefore, a major challenge in understanding disease susceptibility is to determine how non-coding DNA variants disrupt CREs. A further challenge is that DNA variants, such as expression quantitative-trait loci (eQTLs), often have effects in a single cell type^3^ or stimulation condition^5,6^. Such studies highlight the need for experimental approaches to characterize the impact and mechanisms of non-coding DNA variants on CRE function in a cell state-specific manner.

Current high-throughput approaches to study the molecular mechanisms by which ncSNPs alter gene expression are based primarily on computational predictions of TF binding^4,7,8^ or on allelic imbalance in genomic assays of TF binding and chromatin state^7,9–13^. However, these approaches have various limitations. Genomic assays based on allelic imbalance are impractical as a general approach to study candidate ncSNPs because each DNA variant must be present in the assayed cells and each experiment can examine only a single TF or chromatin feature. Computational approaches that use position-weight matrix (PWM) models to assess the impact of ncSNPs on TF binding offer a parallelizable approach, but can predict altered TF binding for only a fraction of ncSNPs^4,14^. Additionally, PWM-based approaches do not account for changes in TF activity, such as TF nuclear localization or interactions with COFs, that occur in response to cell-state changes and are known to affect ncSNP function^5,6^.

To address these challenges in ncSNP annotation, we have developed CASCADE (<u>C</u>omprehensive <u>AS</u>sessment of <u>C</u>omplex <u>A</u>ssembly at <u>D</u>NA <u>E</u>lements) – a protein-binding microarray (PBM)-based high-throughput approach to profile the DNA binding of TF-COF complexes from cell nuclear extracts. PBMs are double-stranded DNA microarrays that allow protein-DNA binding to be assayed to thousands of DNA sequences^15,16^. Recently, we developed the nuclear extract protein-binding microarray (nextPBM) approach^17^ to study DNA binding of TFs present in nuclear extracts. However, TFs function by recruiting COFs, which subsequently alter gene expression through diverse mechanisms such as histone modification or chromatin remodeling (Figure 1a)^18^. To directly interrogate TF-COF complexes, in CASCADE we extend nextPBM to profile *recruitment* of COFs to DNA variants using nuclear extracts

**Figure 1.**
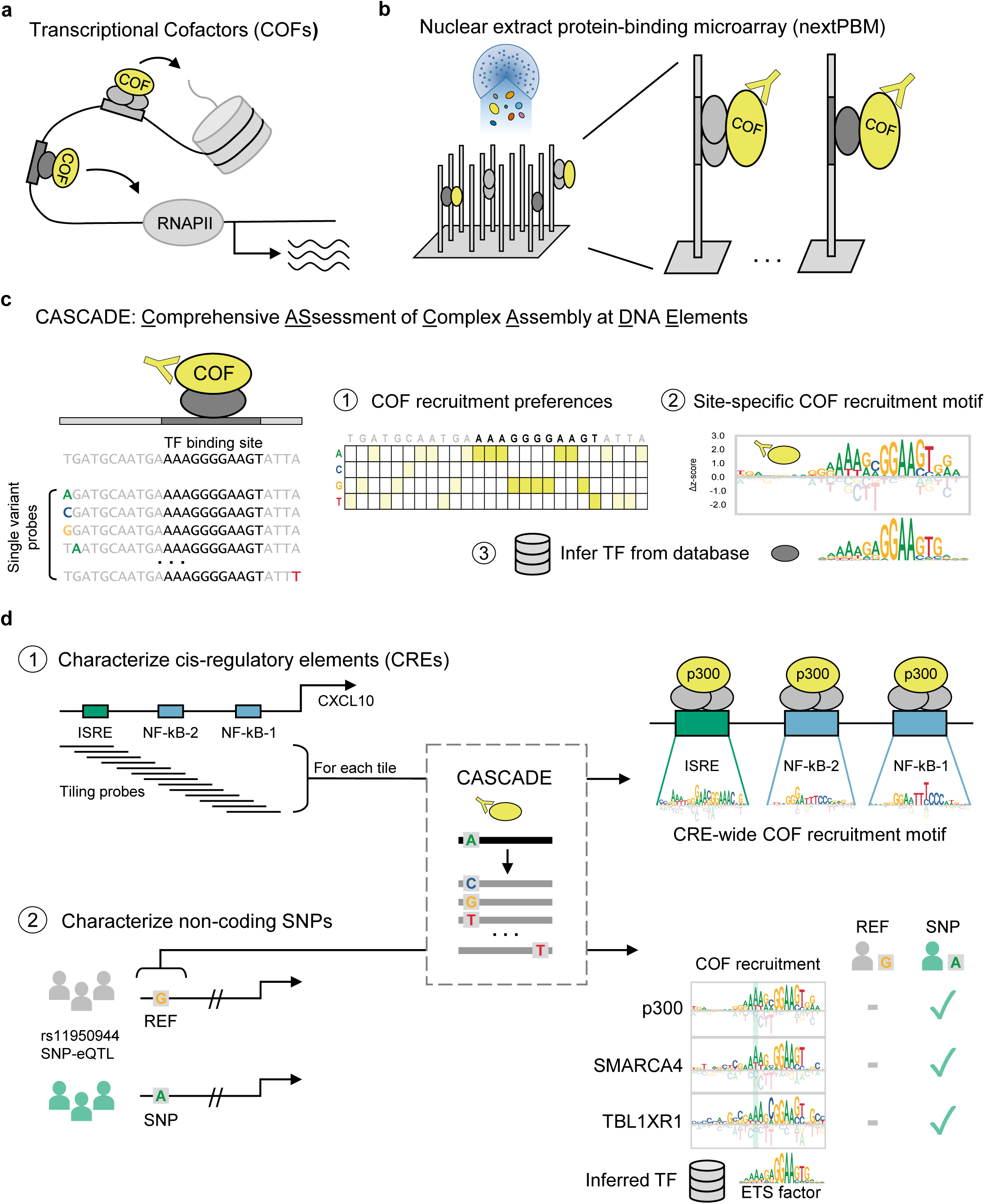
Comprehensive Assessment of Complex Assembly at DNA Elements (CASCADE) approach and applications. (**a**) COFs affect transcription and chromatin state. (**b**) COF recruitment to DNA is assayed by nextPBM. (**c**) COF recruitment is assayed to a ‘seed’ probe (e.g., genomic-derived TF binding site sequence) and all single variant probes. (1) COF recruitment to single variant probes yields nucleotide preferences along DNA sequence. (2) Preferences transformed to COF recruitment motif (i.e., a logo). (3) Motif matched to TF motif databases to infer TF identity. (**d**) Overview of CASCADE applications. CASCADE can be applied (1) cis-regulatory elements (CREs) or (2) reference (REF) / non-coding single-nucleotide variant (ncSNP) pairs. For CRES, tiling probes are used to span the genomic region, and COF motifs for each tiling probe are integrated into a CRE-wide COF motif. For ncSNP/REF pairs, COF motifs are determined for both and compared. ISRE: interferon stimulated response element, NF-κB: Nuclear Factor kappa light chain enhancer of activated B cells.

(Figure 1b). As many COFs, such as the acetyltransferase EP300/CBP, interact broadly with multiple TFs^19–21^, we can assay many TF-COF complexes in a parallel manner by profiling recruitment of a single COF, without requiring previous knowledge of the TFs involved. Critically, by assaying COF recruitment to single-nucleotide variants of a DNA sequence, we can determine a COF *recruitment* motif whose specificity allows us to infer the identity of the TF (or TF family) by comparison against TF motif databases (Figure 1c). Therefore, conceptually, by profiling the recruitment of a limited set of COFs we can characterize the DNA binding of a much larger set of TF-COF complexes. Here, we demonstrate that CASCADE can be used to profile the DNA-sequence dependence of TF-COF complex binding to CREs or ncSNPs in a cell-state specific manner (Figure 1d), providing a high-throughput approach to address the biophysical impact of non-coding DNA variants on gene regulatory complexes.

### Application of CASCADE to CREs

To demonstrate the use of CASCADE to characterize CREs, we profiled the recruitment of the COF EP300, hereafter p300, to a promoter segment of the chemokine gene *CXCL10* in resting and lipopolysaccharide (LPS)-stimulated human THP-1 macrophages. CXCL10 is important for mediating the inflammatory response by promoting activation and recruitment of several types of immune cells, such as monocytes. The expression of *CXCL10* is often dysregulated in autoimmune diseases and has been implicated in cancer pathogenesis^22,23^. In LPS-induced activation of *CXCL10* in macrophages, three separate TF binding sites in the promoter are required for full activation, two NF-κB binding sites and an interferon-sensitive response element (ISRE)^24,25^ (Figure 2a), providing a test case for our CASCADE approach. p300 is a broadly acting acetyltransferase that is recruited by diverse TFs, including both NF-κB and IRF3 that function at the *CXCL10* promoter^19,24,25^.

**Figure 2.**
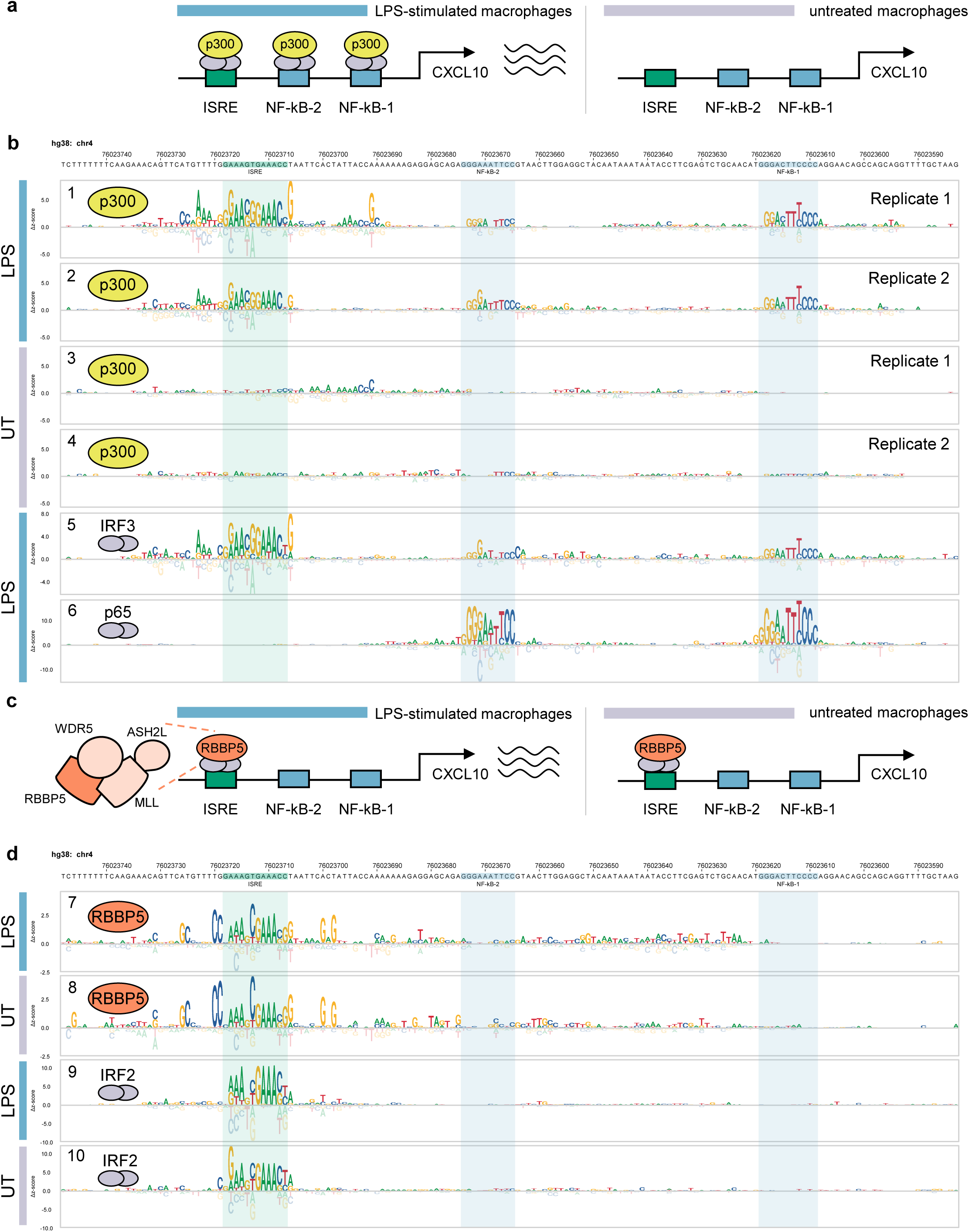
CASCADE-based characterization of COF recruitment to the *CXCL10* promoter. (**a**) Schematic of LPS-inducible recruitment of p300 to *CXCL10* promoter in macrophages. (**b**) CRE-wide p300 recruitment motif and TFs IRF3 and p65/RELA across *CXCL10* promoter. Experiments using extracts from LPS-stimulated or untreated (UT) macrophages are indicated with colored bars. p300 motifs are shown for biological replicate experiments (Replicate 1 and 2). (**c**) Schematic of condition-independent recruitment of RBBP5 to *CXCL10* promoter. (**d**) CRE-wide motifs for COF RBBP5 and TF IRF2 across the *CXCL10* promoter segment. Experimental conditions as in (**b**).

To query p300 recruitment across the *CXCL10* promoter segment (166 bp), we assayed recruitment to 29 tiling probes (each 26 bp long) generated at 5 bp intervals across the target promoter region (Figure 1d, see Methods, Supplementary Data 1). For each tiling probe on our microarray, we also included all single variant (SV) probes to allow a COF recruitment motif to be determined every 5 bp (Figure 1c, 1d). A CRE-wide p300 recruitment motif was then generated for each experimental condition by integrating these individual motifs across their overlapping positions (Figure 2b, tracks 1-4, see Methods).

Our CRE-wide recruitment motif revealed p300 recruitment to the three previously characterized TF binding sites occurred in an LPS-inducible manner (Figure 2b, tracks 1-4). These results are consistent with previous studies that demonstrated the LPS-inducible binding of IRF3 and NF-κB to the *CXCL10* promoter^25–29^. To infer the identity of the TFs involved, we compared the p300 recruitment motifs to a database of previously characterized TF binding motifs (see Methods) and identified IRF3 and NF-κB as high-scoring matches (Supplementary Figure 1a, track 1, Supplementary Figure 1b, track 2). To confirm the binding of NF-κB and IRF3 at these sites, we also performed CASCADE experiments directly for the TFs RELA (the p65 subunit of NF-κB) and IRF3, using antibodies against the TFs instead of p300. p65 bound specifically to the previously characterized NF-κB sites and exhibited the expected DNA binding site specificity (Figure 2b, track 6, Supplementary Figure 2, track 14). IRF3 bound specifically to the ISRE^28,30^ and weakly to the two NF-κB sites, which is consistent with the indirect tethering of IRF3 by NF-κB previously reported in LPS-stimulated macrophages (Figure 2b, track 5)^31,32^. Critically, the binding motifs determined for IRF3 (Figure 2b, track 5) and p65 (Figure 2b, track 6) agree strongly with those for p300 (Figure 2b, tracks 1-2) demonstrating that COF recruitment motifs can accurately capture the binding motifs for the underlying TFs.

To determine whether additional COFs with different effector functions are also recruited to the *CXCL10* promoter segment, we profiled the recruitment of RBBP5, a core subunit of the MLL histone lysine methyltransferase complex (Figure 2c, Supplementary Data 1). Unlike the LPS-inducible recruitment of p300, RBBP5 is constitutively recruited to the *CXCL10* promoter sequences at comparable levels in the presence or absence of LPS (Figure 2d, tracks 7-8). RBBP5 is recruited only to the ISRE element, and not the NF-κB sites, demonstrating a different recruitment preference than p300. However, as IRF3 binding to the ISRE is LPS-induced (Figure 2b, track 5, Supplementary Figure 2, track 13), our data suggests recruitment of RBBP5 to this site is dependent on a different TF. Furthermore, the COF recruitment motifs for p300 and RBBP5 at the ISRE site exhibit clear differences in nucleotide preference (e.g., RBBP5 prefers a 5’-AAANC<u>GAAA</u>-3’ consensus whereas p300 prefers a 5’-GAACG<u>GAAA</u>-3’ consensus; Figure 2b, tracks 1-2, Figure 2d, tracks 7-8). Comparing the RBBP5 recruitment motifs against a TF motif database (see Methods), we identified IRF2 as a high-scoring match (Supplementary Figure 1c, track 7, Supplementary Figure 1d, track 8). IRF2, and the related IRF8, are both constitutively expressed in THP-1 macrophages, which would support the LPS-independent RBBP5 recruitment. CASCADE analysis of both IRF2 and IRF8 yielded CRE-wide motifs that closely matched those obtained for RBBP5 (Figure 2d, tracks 9-10, Supplementary Figure 2, tracks 11-12). These results show that applying CASCADE to different COFs can reveal TF-COF complexes with distinct compositions and DNA-binding specificities.

### Application of CASCADE to ncSNPs

To investigate the extent to which ncSNPs function by perturbing TF-COF complex binding, we used nextPBM/CASCADE approaches to screen ncSNPs for altered COF recruitment. To increase the number of ncSNPs that we could screen, we developed a hierarchical two-step approach to identify and characterize SNPs that affect binding of TF-COF complexes (Figure 3a). In step one, COF recruitment to pairs of reference and SNP alleles is screened in order to identify variants that lead to significant differential COF recruitment (Figure 3, step 1). In step two, to infer the identity of the TFs involved at each SNP locus, a second microarray is used to perform a CASCADE-based analysis for these significant loci (Figure 3, step 2). The COF recruitment motifs generated for each SNP locus can then be compared to TF motif databases to infer the identity of the TF family and to provide additional context for assessing the impact of each SNP.

**Figure 3.**
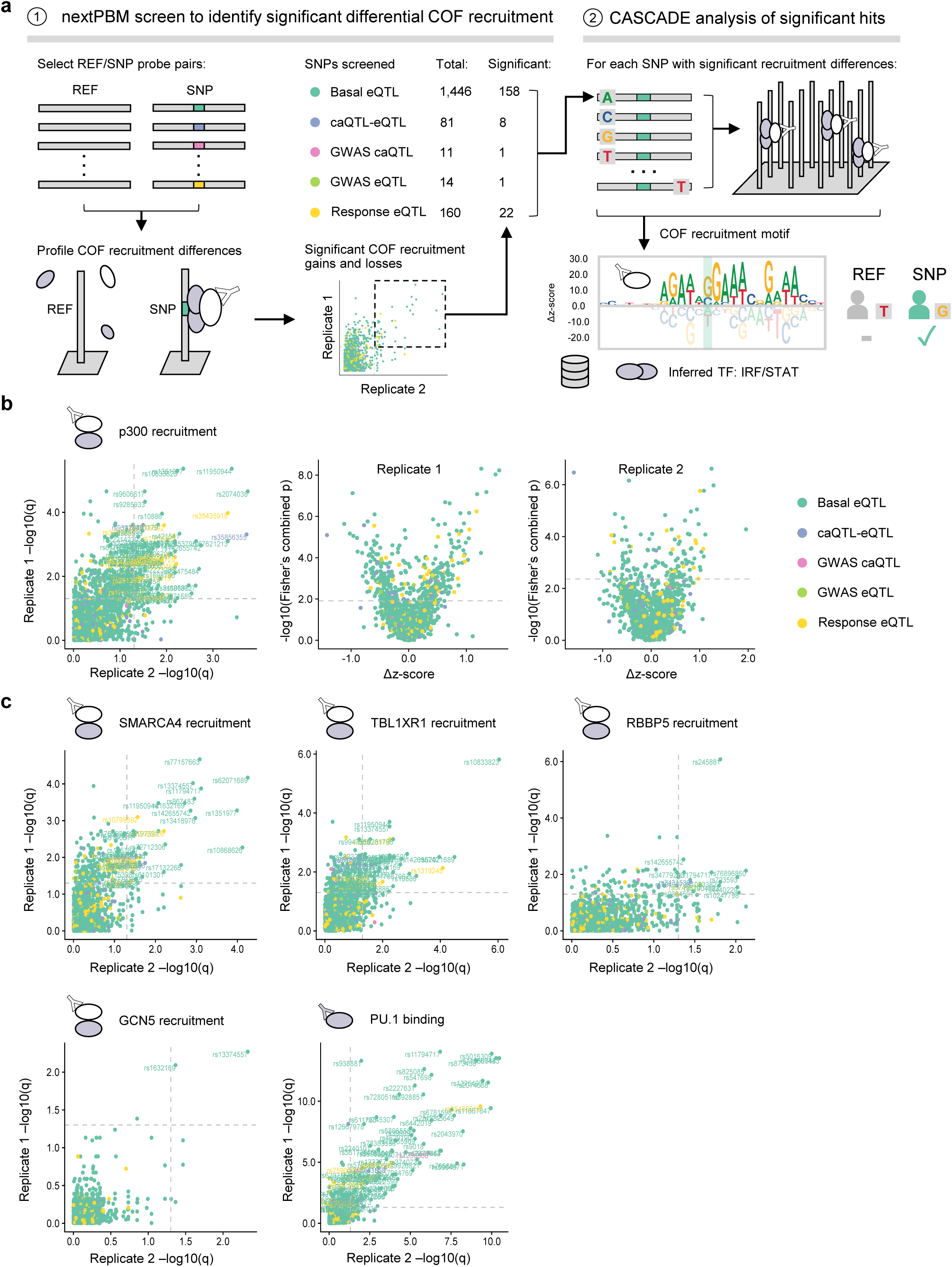
CASCADE-based analysis of SNP-QTLs in human macrophages. (**a**) Overview of 2-step, CASCADE-based approach to characterize 1,712 SNPs/QTLs. (1) Step 1: screen for differential COF recruitment to SNP-QTL/REF probe pairs. Number of probe pairs in each QTL class for which significant COF recruitment was identified in at least one experiment. (2) Step 2: CASCADE-based motifs are generated for SNPs identified as significantly bound. COF motifs are compared against TF-motif databases to infer TF identity. (**b**) Comparison of p300 differential recruitment across biological replicates. Comparison of Q-values for replicates is shown (left). Comparison of differential nextPBM z-scores for SNP/REF pairs against Q-values is shown for replicate experiments. QTL class for each SNP is indicated. (**c**) Comparison of differential COF recruitment across biological replicates is shown for candidate COFs and the TF PU.1. eQTL: expression quantitative trait loci, caQTL: chromatin accessibility quantitative trait loci.

We used this two-step approach to profile COF recruitment to 1,712 SNPs associated with gene expression (eQTLs) and chromatin accessibility (caQTLs) changes in myeloid cells^3,5,6^ (Figure 3a, Supplementary Data 2). We performed our analysis with nuclear extracts from THP-1 macrophages stimulated with IFN-γ and LPS (see Methods). To assess the impact of SNPs on different cellular functions, we profiled recruitment of five COFs from different functional categories: p300, a histone acetyltransferase; SMARCA4/BRG1, a subunit of the SWI/SNF chromatin remodeling complex; TBL1XR1, a subunit of the nuclear receptor corepressor (NCoR) complex; RBBP5, a subunit of the MLL histone lysine methyltransferase complex; and GCN5, a histone acetyltransferase. In addition to these COFs, we screened for differential binding of the TF PU.1 due to its known role in establishing the myeloid enhancer landscape and the previously demonstrated prevalence of the PU.1 binding motif at macrophage SNP-QTLs3,33,34.

Our step-one screen identified 164 total SNPs that reproducibly altered the recruitment of at least one of the tested COFs (Figure 3b, 3c), representing 9.6% of the sites examined.

With the exception of the GWAS caQTL category, comparable proportions of the SNP-QTL categories tested reproducibly altered COF recruitment: 136 basal eQTLs (9.4%), 7 caQTL-eQTLs (8.6%), 1 GWAS eQTL (7.1%) and 20 response eQTLs (12.5%). Profiling the TF PU.1, we also observed widespread differential PU.1 binding at 95 SNP-QTLs (Figure 3c) including 23 that coincided with the differential recruitment of at least one of the COFs screened.

By examining the direction of the differential recruitment, we identified SNPs that caused gain or loss of TF-COF binding (Figure 3b, Supplementary Figure 3). For example, our screen identified 63 SNP alleles that led to statistically significant gain of p300 recruitment (Figure 3b, rightmost two panels, positive Δz-score) and 35 SNP alleles that led to a significant loss relative to the reference allele (Figure 3b, rightmost two panels, negative Δz-score). In total, across all COFs and TFs screened, we observed differential recruitment/binding at 243 of the 1,712 SNP-QTLs (14.2%) with 134 gains, 108 losses, and one SNP demonstrating both. Of note, for each SNP exhibiting significant reproducible differential recruitment of more than one COF (40 total), the direction of the effect, either gain or loss, was consistent across each COF. These results demonstrate that our nextPBM COF-based approach can be used to reproducibly screen broad classes of ncSNPs for both gains or losses of TF-COF complex binding.

For step two of our SNP analysis we used CASCADE to determine COF recruitment motifs at select loci. These motifs allow us to infer the identity of the TFs mediating differential COF recruitment at each locus (Figure 3a, step 2). We selected 158 basal eQTLs, 8 caQTL-eQTLs, 1 GWAS caQTL, 1 GWAS eQTL, and 22 response eQTLs, as these loci showed significant differential recruitment of one or more of the regulators screened (see Methods, Supplementary Data 3). To determine our COF recruitment motifs, we profiled the base preferences of the local genomic region (26 bp) centered at each of these SNP-QTLs. Consistent with our observed differential PU.1 binding, the COF recruitment motifs for many loci matched ETS-type binding motifs (Figure 4). COF motifs were also identified that matched TBX/KLF/EGR zinc finger motifs, IRF/STAT motifs, and two motifs that did not match a known TF motif at a relaxed stringency threshold (Figure 4, see Methods). Comparing the recruitment motifs generated at a given SNP locus, we found the motif base preferences and alignment were consistent across COF and PU.1 experiments, confirming a common underlying TF-COF complex. Examining SNPs specifically affecting ETS motifs, we found that SNPs can impact different positions along the ETS motif, including both the variable 5’ flanking region (rs11940944, rs72755909, rs2526718) and the core ETS 5’-GGAA-3’ element (rs873458, rs1250568). These results highlight that COF recruitment motifs can provide a means to understand the biophysical mechanism for a SNP-QTL.

**Figure 4.**
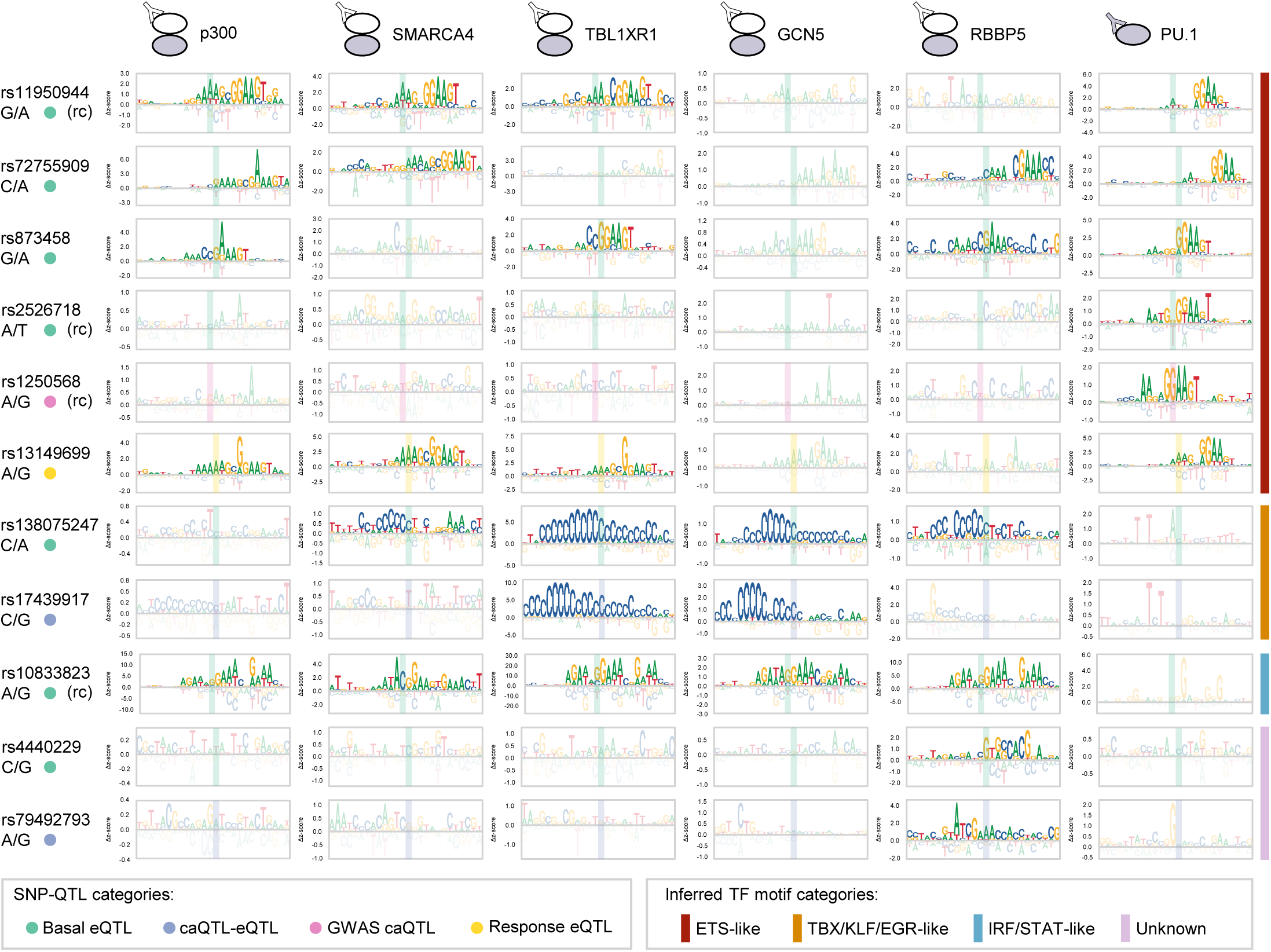
CASCADE-determined motifs at SNP loci. COF recruitment motifs for p300, SMARCA4, TBL1XR1, GCN5, and RBBP5 are shown for 10 SNP-QTL loci. PU.1 binding motifs at each locus are also shown. Position of the SNP location within each motif is shown with a shaded rectangle. QTL type of each SNP is indicated (left-hand side, colored dots). Only sites that met an imposed seed z-score threshold were plotted (see Methods). Corresponding reference and SNP are shown beneath each rsID. (-) denotes a site plotted as its reverse complement relative to the reference strand. For these sites, the reference and SNP alleles are also indicated as their complementary nucleotides.

We highlight two gain-of-recruitment SNP-eQTLs identified in our screen to demonstrate how CASCADE can be used to generate mechanistic models of ncSNPs. Our analysis for rs11950944 (G/A), a basal SNP-eQTL in myeloid cells^5^, found that p300 (z-score: 2.36), SMARCA4 (z-score: 2.99), and TBL1XR1 (z-score: 2.61) are recruited to the SNP allele but are either not recruited or are below our detection threshold for the reference allele (p300: z-score: - 0.13, SMARCA4: z-score: -0.38, TBL1XR1: z-score: 0.37) (Figure 5a left, Supplementary Data 3). The COF recruitment motifs for all three COFs matched significantly with ETS-factor motifs (Figure 5a, right). Consistent with our motif-based inferences, the ETS factor PU.1 preferentially bound the SNP allele (z-score: 5.99) though it could also be detected at the reference allele (z-score: 4.04). These results suggest a model where the SNP allele enhances the DNA binding of an ETS-family TF, possibly PU.1, which leads to enhanced recruitment of these COFs (Figure 5c). We note that enhanced binding of PU.1 at DNA variants in murine myeloid cells has been previously shown to correlate with increased local histone modifications characteristic of primed and active regulatory elements as well as with increased transcriptional output^34^.

**Figure 5.**
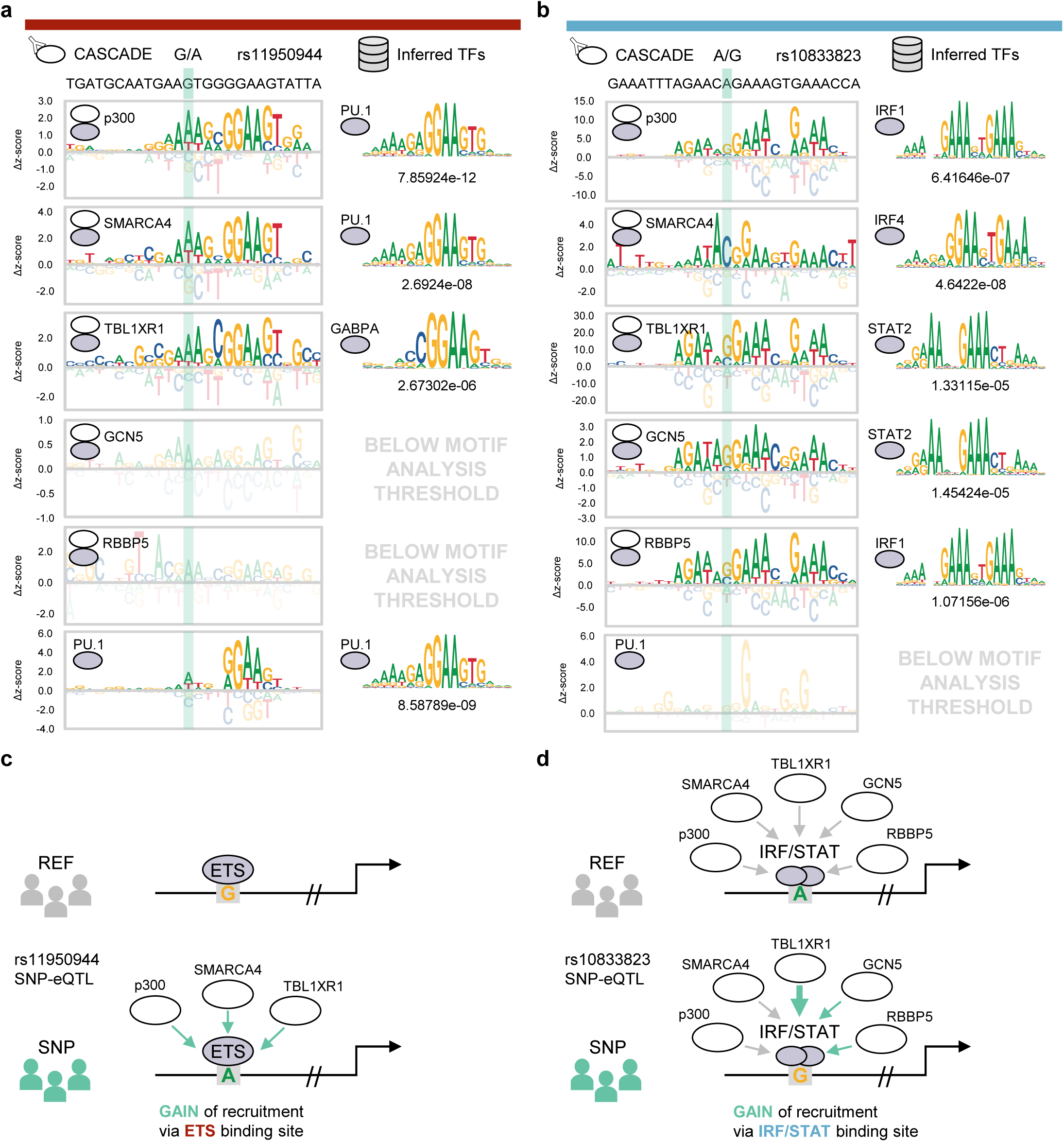
Constructing models with CASCADE for SNP-eQTLs. (**a**) Left column: CASCADE-determined COF recruitment motifs for p300, SMARCA4, TBL1XR1, GCN5, and RBBP5 at the local genomic region surrounding rs11950944. PU.1 binding motif is also shown. Right column: TF binding motif with the strongest association to each corresponding CASCADE COF recruitment motif. Statistical significance (p-value) for TF matching is shown below each TF motif (see Methods). Position of the SNP location within each motif is shown in the shaded area. QTL type and inferred TF category are indicated by the same color scheme as in Figure 4. (**b**) Same as in (**a**) but for the local genomic region surrounding rs10833823. Only sites that met an imposed z-score threshold were plotted and used for motif analysis (see Methods). (**c**) Integrative model for COF recruitment changes at SNP-eQTL rs11950944. (**b**) Same as in (**c**) but for SNP-eQTL rs10833823.

Our analysis for a second basal SNP-eQTL rs10833823 (A/G) in myeloid cells^5^ identified a different scenario in which the entire panel of COFs tested were recruited to the reference allele, but the SNP allele caused significantly higher recruitment for three of the COFs: TBL1XR1 (z-scores: WT = 9.71, SNP = 28.49), GCN5 (z-scores: WT = 1.54 to SNP = 3.36), and RBBP5 (z-scores: WT = 10.56 to SNP = 15.82) (Figure 5b left, Supplementary Data 3). The COF recruitment motifs for all COFs matched GA-rich IRF/STAT-family motifs (Figure 5b, right), and consistent with our inference of recruitment by IRF/STAT-type TFs, we did not observe PU.1 binding at this site (Figure 5b, left). Notably, while the SNP allele enhanced COF recruitment in our assay it occurred at a low-information position in the IRF/STAT motifs that did not appreciably affect the PWM scores for these TFs. Therefore, a computational screen for this SNP would not predict appreciable changes in TF binding and would be missed. Our results for this SNP suggest that the SNP allele does not affect TF binding but can alter the recruitment of COFs (Figure 5d), possibly by a mechanism involving DNA-based allostery^35,36^. These results demonstrate how the CASCADE approach, based on COF-recruitment profiling, can generate biophysical, mechanistic models for how ncSNPs can alter the binding of TF-based regulatory complexes.

## Discussion

Characterizing the effects of DNA variants, such as ncSNPs, on gene regulatory complexes is a challenge in our efforts to explain the genetic contributions to human disease. A bottleneck in the field is that studies identifying the mechanisms by which ncSNPs function greatly lag studies identifying ncSNPs associated with traits or diseases^2^. To address this need for high-throughput approaches to characterize ncSNPs, we developed CASCADE as a high-throughput, customizable platform for profiling the impact of DNA variants on TF-COF complexes. By measuring the DNA recruitment of broadly interacting COFs (i.e., that form complexes with many TFs), this approach can assay multiple TF-COF complexes in a multiplexed manner. Furthermore, as CASCADE queries the binding of TF-COF complexes, as opposed to just TFs, it can suggest a link between DNA variants and the biological functions mediated by each COF. In this work, we have applied CASCADE to the study of ncSNPs, but the approach can be customized to study any non-coding DNA variants, such as rare variants associated with disease or somatic mutations associated with cancer. We envision that using CASCADE in conjunction with other high-throughput, cell-based methods, such as massively-parallel reporter assays (MPRAs) that assess gene expression^37–39^, will provide exciting new approaches to characterize function and mechanism of DNA variants at a genomic scale.

As disease-associated ncSNPs often reside within CREs^3,4,6,40^ the characterization of ncSNPs is directly related to the problem of delineating the mechanisms of CREs. Here, we demonstrate that CASCADE can be applied to this fundamental problem and can be used to identify TF binding sites within CREs and the TF-COF complexes that bind to these sites under different cellular conditions. Using CASCADE to characterize an LPS-inducible segment of the *CXCL10* promoter, we identified the three previously validated NF-κB and IRF sites involved and TF-COF complexes bound to each individual site. In this work, we profiled a limited set of COFs, but the approach can be applied to other COFs where native antibodies are available, or COFs have been affinity tagged. We also demonstrated that we can identify site-specific recruitment of COFs that are annotated as subunits of larger, multi-protein COF complexes (e.g., RBBP5, Figure 2c). Currently it is unclear the extent to which these multi-protein COF complexes are assembled on our microarrays, or whether we are assaying the recruitment of smaller sub-complexes or even single COFs (i.e., binary TF-COF interactions). Future studies will address the extent to which recruitment of larger COF complexes is being assayed and how CASCADE-identified TF-COF interactions reflect interactions critical for CRE function *in vivo*. Finally, we note that CASCADE provides the first, high-throughput approach to establish the link between individual CRE binding sites, TFs, and COFs. We anticipate that CASCADE will allow for renewed examination of the role COFs play in the cis-regulatory logic that governs CRE function, which has primarily focused on TFs and binding sites alone.

High-throughput methods for studying TF-DNA binding (e.g., MITOMI, SMiLE-seq, CSI, PBM, SELEX-seq, ATI, nextPBM, etc.)^15–17,41–47^ have had a tremendous impact on our understanding of TF function and genome-scale analysis of gene regulation, leading to large databases of widely-used TF binding models^48–51^. However, these methods have not been applied to the study of TF-COF complexes. Approaches such as ATI^47^ and nextPBMs^17^ have been used to study TF-DNA binding in a more native cellular context by using TFs directly from cell nuclear extracts instead of using purified TF samples. However, COF recruitment has not been examined using these approaches. Here, we demonstrate with CASCADE that COF recruitment, and by extension the assembly of TF-COF complexes, can be directly profiled in a high-throughput manner using cell nuclear extracts. Analogous to TF binding motifs, we introduce the concept of a COF *recruitment* motif that represents the DNA sequence-specificity of COF recruitment. We demonstrate that COF motifs can be used to infer the identity of the TF (or TF family) recruiting a COF to a particular DNA sequence. We note that this approach is different from COF ChIP-seq, which identifies the genomic loci to which a COF is recruited, but does not identify the TFs involved at individual loci, nor the DNA-sequence dependence of COF recruitment at single-nucleotide resolution. Given that COFs can be recruited by multiple TFs, by assaying recruitment of a single COF we are able to profile numerous TF-COF complexes in parallel. CASCADE offers a conceptually new high-throughput approach to study gene regulatory complexes. We anticipate that the ability to assay COF recruitment afforded by CASCADE will provide a deeper general understanding of how DNA sequence and regulatory complexes control gene expression and cellular responses.

The CASCADE approach introduced here is a scalable, customizable platform to study TF-COF complexes and the impact of DNA variants on these gene regulatory complexes. In this study, we demonstrate its application to the functional characterization of CREs and ncSNPs. However, it can be customized and applied to many other types of DNA variants and elements in a cell-specific manner, such as mutations in different cancers or synthetic regulatory elements designed to drive a cell-specific response. We show the application of CASCADE to nuclear extracts from a human macrophages cell line, but conceptually the approach can be used with nuclear extracts from any cell or tissue type. Finally, the ability to profile COF recruitment to DNA sites provides an opportunity to link DNA variants with therapeutic intervention. COFs are often enzymatic (e.g., methyltransferases, histone deacetylases, etc.) and therapeutic inhibitors for many COFs are available^52,53^. Identifying the TF-COF complexes whose binding site is created by a DNA variant may allow for the identification of therapeutic antagonists to counteract their effects. Future studies applying CASCADE in these diverse scenarios should help to develop the approach and provide insights into the roles of TF-COF complexes in cell signaling and disease.

## Acknowledgments

National Institutes of Health (NIH) R01 grant (R01A116829 to T.S.). We thank Juan Fuxman Bass, Thomas Gilmore, Andrew Emili, and David J. Waxman for discussions related to this work.

## Author contributions

D.B, H.H, and T.S. jointly conceived of the study. D.B. developed the software to design, analyze, and visualize CASCADE nextPBM experiments. H. H. performed the experimental work. J.K., A.P., N.M., R.Z., and Y.O. supported H.H. with validation of the experimental technique. D.B., H.H., and T.S. analyzed and interpreted the experimental results. T.S. oversaw the experimental work and development of the computational methods. D.B., H.H., and T.S. wrote the manuscript.

## Methods

### Cell Culture

THP-1 cells, a human monocyte cell line, were obtained from ATCC (TIB-202). The cells were grown in suspension in RPMI 1640 Glutamax media (Thermofisher Scientific, Catalogue #72400120) with 10% heat-inactivated fetal bovine serum (Thermofisher Scientific, Catalogue #11360070) and 1mM sodium pyruvate (Thermofisher Scientific, Catalogue #16140071). T175 (Thermofisher Scientific, Catalogue #132903) non-treated flasks were used when culturing THP-1 cells for experiments. Cells were grown in 50mL of media when being cultured in T175 flasks.

To differentiate THP-1 cells into adherent macrophages, cells were grown to a density of 8.0 × 10^5^ cells/mL and treated with 25ng/mL Phorbol 12-Myristate 13-Acetate (PMA) (Sigma-Aldrich, Catalogue #P8139) for 4 days. Following the 4 days of PMA treatment, the media was replaced with fresh RPMI media with 10% heat-inactivated fetal bovine serum and 1mM sodium pyruvate. The cells rested for two days in the fresh media before being stimulated with various reagents.

THP-1 cells differentiated with PMA were treated with either Lipopolysaccharide (LPS) (Sigma-Aldrich, L3024) or Interferon gamma (IFN-γ) (Thermofisher Scientific, Catalogue #PHC4031) in combination with LPS. PMA treated THP-1 cells were treated with 1ug/mL of LPS for 45 min or with 100ng/mL IFN-γ for 2 h followed by 1ug/mL LPS for 1 h. For each condition, nuclear lysates were harvested. For all nuclear lysates assayed using PBM experiments, the expression levels of COFs and TFs profiled with CASCADE were confirmed by western blotting (Supplementary Figure 4).

### nextPBM experimental methods

The nuclear extract protocols are as previously described^17^. Changes to the previously published protocols are detailed. To harvest nuclear extracts from THP-1 cells, the media was aspirated off and the cells were washed once with 1X PBS (Thermofisher Scientific, Catalogue #100010049). Once the 1X PBS used to wash the cells was aspirated off, enough 1X PBS was mixed with 0.1mM Protease Inhibitor (Sigma-Aldrich, Catalogue #P8340) to cover the cells was added to each flask. A cell scraper was then used to dislodge the cells from the flask. The cells were collected in a falcon tube and placed on ice. To pellet the cells, the cell volume was centrifuged at 500xg for 5 min at 4°C. Once the cells were pelleted, the supernatant was aspirated off. The pellet was resuspended in Buffer A and incubated for 10 min on ice (10mM HEPES, pH 7.9, 1.5mM MgCl, 10mM KCl, 0.1mM Protease Inhibitor, Phosphatase Inhibitor (Santa-Cruz Biotechnology, Catalogue #sc-45044), 0.5mM DTT (Sigma-Aldrich, Catalogue #4315)) to lyse the plasma membrane. After the 10 min incubation, a final concentration of 0.1% Igepal detergent was added to the cell and Buffer A mixture and vortexed for 10 sec. To separate the cytosolic fraction from the isolated nuclei, the sample was centrifuged at 500xg for 5 min at 4°C. The cytosolic fraction was collected into a separate microcentrifuge tube. The pelleted nuclei were then resuspended in Buffer C (20mM HEPES, pH 7.9, 25% glycerol, 1.5mM MgCl, 0.2mM EDTA, 0.1mM Protease Inhibitor, Phosphatase Inhibitor, 0.5mM DTT, and 420mM NaCl) and then vortexed for 30 sec. The nuclei were incubated in Buffer C while mixing at 4°C. To separate the nuclear extract from the nuclear debris, the mixture was centrifuged at 21,000xg for 20 min at 4°C. The nuclear extract was collected in a separate microcentrifuge tube and flash frozen using liquid nitrogen. Nuclear extracts were stored at -80°C until used for experiments.

Microarray DNA double stranding and PBM protocols are as previously described^15–17^. Any changes to the previously published protocols are detailed. Double-stranded microarrays were pre-wetted in HBS (20mM HEPES, 150mM NaCl) containing 0.01% Triton X-100 for 5 min and then de-wetted in an HBS bath. Next the array was incubated with nuclear extract for 1 h in the dark in a binding reaction buffer (20mM HEPES, pH 7.9, 100mM NaCl, 1mM DTT, 0.2mg/mL BSA, 0.02% Triton X-100, 0.4mg/mL salmon testes DNA (Sigma-Aldrich, Catalogue #D7656)). The array was then rinsed in an HBS bath containing 0.1% Tween-20 and subsequently de-wetted in an HBS bath. After the protein incubation, the array was incubated for 20 min in the dark with 20ug/mL primary antibody for the TF or COF of interest (Supplementary Table 1). The primary antibody was diluted in 2% milk in HBS. After the primary antibody incubation, the array was first rinsed in an HBS bath containing 0.1% Tween-20 and then de-wetted in an HBS bath. Microarrays were then incubated with 10ug/mL of either alexa488 or alexa647 conjugated secondary antibody (see Supplementary Table 1) for 20 min in the dark. The secondary antibody was diluted in 2% milk in HBS. Excess antibody was removed by washing the array twice for 3 min in 0.05% Tween-20 in HBS and once for 2 min in HBS in coplin jars as described above. After the washes, the array was de-wetted in an HBS bath. Microarrays were scanned with a GenePix 4400A scanner and fluorescence was quantified using GenePix Pro 7.2. Exported fluorescence data were normalized with MicroArray LINEar Regression^15^.

### CASCADE microarray designs and analyses

A known LPS-responsive segment of the *CXCL10* promoter (hg38: chr4) from 76023583 to 76023748 was used for the basis of this array design^24^. The genomic region was tiled through using 26-base “target” probe sequences with a 5-base step forward between sequential tiles. In total, 29 of these tile probes were needed to span the LPS-responsive *CXCL10* promoter segment. “Target” sequences corresponding to the genomic locus were obtained from the hg38 genome fasta file included with Bowtie2^54^ using the “fastaFromBed” function from bedtools v2.26.0^55^. For each tile probe and each position along the corresponding 26-base target region, a probe was included in the array design consisting of each possible nucleotide variant (at that position) in order to employ the variant probe analysis approach (see below). A total of 2,291 targets were therefore used to model the *CXCL10* promoter segment (29 tiles + 29 × 3 variant probes x 26 positions). 500 additional 26-base target regions were randomly selected from the hg38 using the bedtools “shuffleBed” function and included in the array design to build a background distribution of fluorescence intensity. Each 26-base target region in the array design was embedded in a larger 60-base PBM probe as follows:

“GCCTAG” 5’ flank – 26-base target region – “CTAG” 3’ flank – “GTCTTGATTCGCTTGACGCTGCTG” double-stranding primer

Each target region was included in its reference (+) orientation as well as the reverse complement (-) orientation. 5 replicate spots of each probe (in each orientation) were included in the final array design. PBM microarray probes, relevant annotation for each, and the experimental results are provided (Supplementary Data 1). The microarrays were purchased from Agilent Technologies Inc. (AMAID: 085605, format: 8×60K).

To design the nextPBM-based screen for differential COF recruitment at ncSNPs, the lead SNPs uncovered in previous studies were included in our high-throughput screen as follows: 1,446 basal eQTLs^5^ (randomly selected from the “classical monocytes” category), 81 caQTL-eQTLs, 11 GWAS caQTLs, 14 GWAS eQTLs, and 160 response eQTLs^3^. Chromosomal coordinates (hg38) for each SNP were obtained using the biomaRt R package from Ensembl^56^. 26-base DNA probe target regions centered at the SNP position (relative to + strand: 13 bases + SNP location + 12 bases) were obtained for each reference (REF) allele using bedtools as above. For each REF allele probe, a probe with the corresponding SNP allele was also included in the design such that each rsID is represented by a pair of REF and SNP probes. 500 background target regions were also included using the same procedure as above. The 26-base target regions were embedded in larger 60-base PBM DNA probes as above. 5 replicates of each probe (in both orientations) were included in the final design. The microarrays were purchased from Agilent Technologies Inc. (AMAID: 085920, format: 8×60K).

Each REF/SNP pair was screened for differential recruitment of p300, SMARCA4, TBL1XR1, RBBP5, and GCN5 as well as differential binding of representative ETS factor PU.1 nextPBM experimental results were preprocessed as above. Z-scores were obtained for each probe as previously described^57^ against the distribution of fluorescence intensities obtained at the set of background probes for a given experiment. For each REF and SNP allele pair in the design, a t-test was used to compare the fluorescence intensity distributions between the 5 REF probes and 5 SNP probes for a given COF/TF assayed. To mitigate the influence of probe orientation-specific effects, t-tests were performed independently for each probe orientation with the p-values combined using Fisher’s method. The Benjamini-Hochberg method was used to adjust the individual p-values for a REF/SNP pair for multiple hypothesis testing. The fluorescence intensity z-score difference for a given REF and SNP allele probe pair (termed Δz-score) was computed by subtracting the mean REF z-score from the mean SNP z-score such that a positive Δz-score represents a gain-of-recruitment introduced by the SNP allele and a negative Δz-score represents a loss. Scatterplots based on the screening results (Figure 3b-c) were plotted using the ggplot2^58^, RColorBrewer^59^, and cowplot^60^ R packages. A full data file including the statistics from the high-throughput differential recruitment screen is included in the supplementary materials (Supplementary Data 2).

Reference and SNP allele pairs exhibiting reproducible significant differential COF recruitment and/or TF binding were selected for this CASCADE array design in order to infer regulators responsible for the differential activity observed. Inclusion criteria was as follows: the difference in recruitment (or binding) of a given COF (or TF) between corresponding REF and SNP allele probes must have obtained an adjusted p-value (q-value) < 0.05 independently in both technical replicates with a concordant direction of effect. Single variant probes for the 26-base target regions (centered at the SNP position – as described above) were generated using the same procedure as above but without the tiling needed to span larger genomic loci such as the *CXCL10* promoter segment used previously. In addition, only 291 background probes were included due to probe number limitations. PBM microarray probes, relevant annotation for each probe, and the experimental results are provided (Supplementary Data 3). The microarrays were purchased from Agilent Technologies Inc. (AMAID: 086248, format: 4×180K).

Motif modeling using single variant (SV) probes was performed as previously described^17,57,61^ for the SNP-QTL sites profiled in detail using CASCADE. For the multi-tile design used to model extended loci such as the LPS-responsive *CXCL10* promoter segment, a weighted mean approach was applied as follows to overlapping positions in order to integrate results across sequential tiles: all variant probes corresponding to a given nucleotide at a given position within the promoter segment were averaged using each probe’s corresponding seed (reference genomic) z-score as a weight. Further, if a given SV probe’s z-score was above 1.645 (above approximately 95% of the fluorescence intensities obtained using background probe distribution - assuming a normal distribution) and the SV probe’s corresponding reference probe z-score was less than or equal to 1.645, the SV probe’s z-score was reset to the reference seed value. This procedure ensured that the SV probe modeling approach was used to characterize true genomic recruitment sites and reduce the influence of COF recruitment sites gained specifically via a non-reference (non-genomic) variant. Sequence logo plots for the COF recruitment and TF binding motifs were generated using the ggseqlogo R package^62^ and arranged using cowplot. The Δz-scores of each nucleotide represent the difference relative to the median z-score obtained across all possible nucleotides at that position and was computed after the weighted averaging procedure described previously. The Δz-score axis limits for the logo tracks (Figure 2, Supplementary Figure 2) were determined using the minimum and maximum Δz-scores obtained for a given COF/TF (across experiments within an array design) to enable comparisons across stimulus conditions assuming matched total protein concentrations across experiments.

### Motif similarity analysis

For CASCADE recruitment motifs obtained at the *CXCL10* locus, to simplify the analysis and reduce the number of comparisons, the promoter segment was first separated into 3 motifs broadly corresponding to each previously characterized TF site (ISRE, NF-κB-2, and NF-κB-1). For CASCADE profiling of the SNP-QTLs, a minimal seed z-score of 1.5 was enforced for motif analysis. Recruitment energy matrices obtained from CASCADE cofactor profiling (fluorescence intensity z-scores) were converted to a probability-based matrix using the Boltzmann distribution as previously described^57^ to be more directly comparable to previous TF binding models:

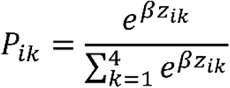

*z*_*ik*_ is the z-score for nucleotide variant *k* at position *i* within the motif window. β transformation parameters for the Boltzmann equation were scaled using the maximum z-score obtained in a given experiment using the following equation in order to account for differences in antibody efficiencies across cofactors:

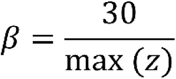

Resulting position-weight matrices were compared against the complete HOCOMOCOv11 database^51^ of transcription factor binding models (771 total) using TOMTOM from the MEME suite^63^ version 5.0.3. Euclidean distance was used as the similarity metric with a relaxed minimal reporting q-value of 0.25 (-dist ed -thresh .25).

## Data availability

The results of all nextPBM/CASCADE array experiments performed here have been deposited in the Gene Expression Omnibus (GEO accession: GSE148945). An R script that implements CASCADE to generate the plots shown in this study using the Supplementary Data files has been made available on Github (https://github.com/Siggers-Lab/CASCADE_paper). All other data is available upon request.

