## Supplementary Information for "Customizable high-throughput platform for profiling cofactor recruitment to DNA to characterize cis-regulatory elements and screen non-coding single-nucleotide polymorphisms"

**Supplementary Data 1. CASCADE nextPBM data for CXCL10 promoter segment.** DNA probe annotations and z-score normalized fluorescence values for CXCL10 CASCADE experiments.

**Supplementary Data 2. Differential COF recruitment summary statistics for SNP-QTL screen.** Statistical significance of COF recruitment differences across paired reference and SNP allele probes for SNP-QTLs screened in this study.

**Supplementary Data 3. CASCADE data for SNP-QTLs with significant differential COF recruitment.** DNA probe annotations and z-score normalized fluorescence values for SNP-QTLs profiled using CASCADE.

**Supplementary Table 1. Antibodies used for experiments.** The antibodies listed were used for DNA pulldown experiments, PBM experiments or both.

| Antibody | Catalog Number | Application |
|---|---|---|
| **Primary Antibodies** | | |
| P300 | ab14984 | PBM Experiment/Western Blot |
| SMARCA4 | sc17796 | PBM Experiment/Western Blot |
| GCN5 | sc-365321x | PBM Experiment/Western Blot |
| RBBP5 | a300-109A | PBM Experiment/Western Blot |
| TBLX1R1 | sc-100908 | PBM Experiment |
| HDAC1 | ab7028 | PBM Experiment |
| P65 | sc-372X | PBM Experiment |
| P65 | sc-8008 | Western Blot |
| IRF8 | sc-6058X | PBM Experiment |
| IRF3 | D83B9 | PBM Experiment |
| IRF2 | sc-374327 | PBM Experiment |
| PU.1 | sc-352X | PBM Experiment |
| **Secondary Antibodies** | | |
| Donkey anti-goat IgG (H+L) Cross-Adsorbed Secondary Antibody, Alexa Fluor 488 | A11055 | PBM Experiment |

| | | |
|---|---|---|
| Goat anti-mouse IgG (H+L) Highly Cross-Adsorbed Secondary Antibody, Alexa Fluor 488 | A11029 | PBM Experiment |
| Goat anti-rabbit IgG (H+L) Highly Cross-Adsorbed Secondary Antibody, Alexa Fluor 488 | A11034 | PBM Experiment |
| Goat anti-mouse IgG (H+L) Highly Cross-Adsorbed Secondary Antibody, Alexa Fluor 647 | A32728 | PBM Experiment |
| Goat anti-rabbit IgG (H+L) Highly Cross-Adsorbed Secondary Antibody, Alexa Fluor 647 | A32733 | PBM Experiment |
| HRP conjugated Goat anti-mouse | G-21234 | Western Blot |
| HRP conjugated Goat anti-rabbit | G-21040 | Western Blot |

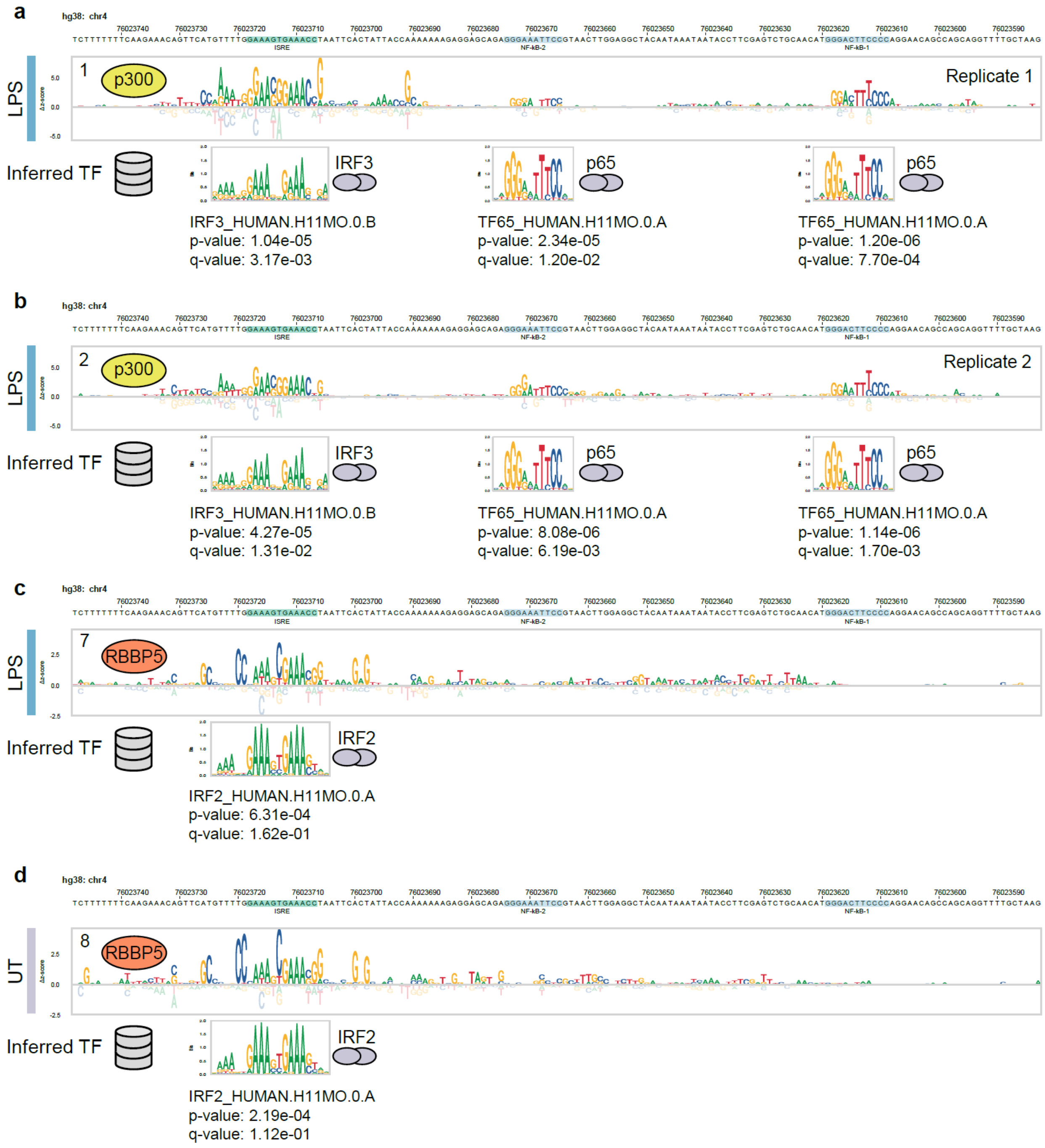


**Supplementary Figure 1. Model-based inference of transcription factors associated with CXCL10 promoter cofactor recruitment motifs.** (**a**) TF motifs matched to p300 recruitment preferences in LPS-stimulated macrophages (Replicate 1). (**b**) TF motifs matched to p300 recruitment preferences in LPS-stimulated macrophages (Replicate 2). (**c**) TF motif matched to

RBBP5 recruitment preferences in LPS-stimulated macrophages (**d**) TF motif matched to RBBP5 recruitment preferences in untreated macrophages. All COF recruitment preference tracks were converted to probability-based models (see Methods) prior to comparison. Similarity comparisons to known TF binding models was performed using TOMTOM and the full HOCOMOCOv11 motif database (771 total motifs – see Methods). COF: transcriptional cofactor, TF: transcription factor, ISRE: Interferon Stimulated Response Element, NF-kB: Nuclear Factor kappa light chain enhancer of activated B cells, LPS: Lipopolysaccharide, UT: Untreated.

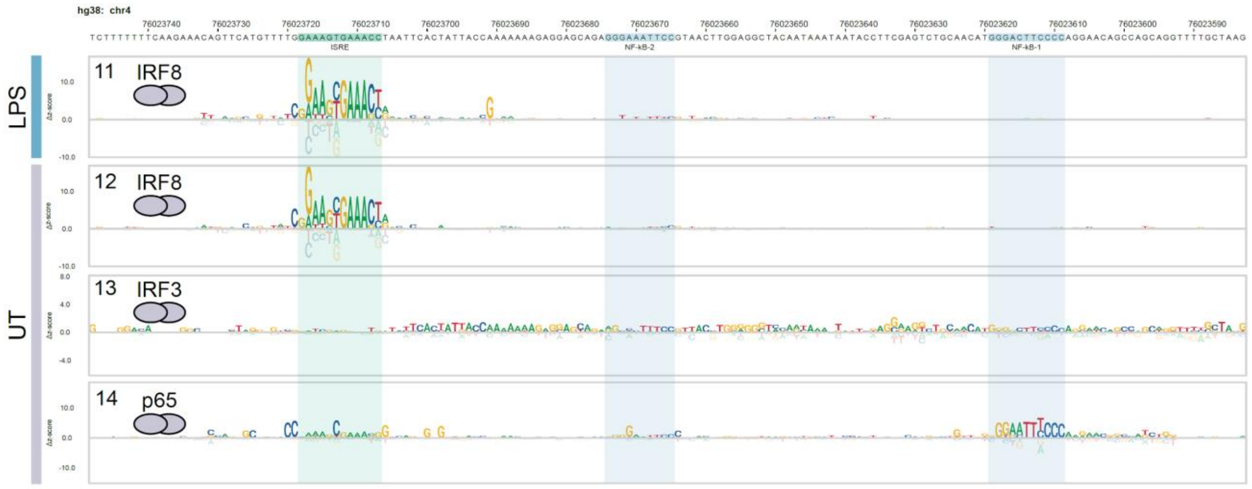


**Supplementary Figure 2. Additional CASCADE-based analyses of transcription factor binding to the CXCL10 promoter segment.** Nucleotide binding preferences of IRF8 to the CXCL10 promoter segment in paired LPS-stimulated (track 11 - continued from Figure 2) and untreated (track 12) macrophages. Binding preferences of IRF3 (track 13) and p65 (track 14) in untreated macrophages. ISRE: Interferon Stimulated Response Element, NF-kB: Nuclear Factor kappa light chain enhancer of activated B cells, LPS: Lipopolysaccharide, UT: Untreated.

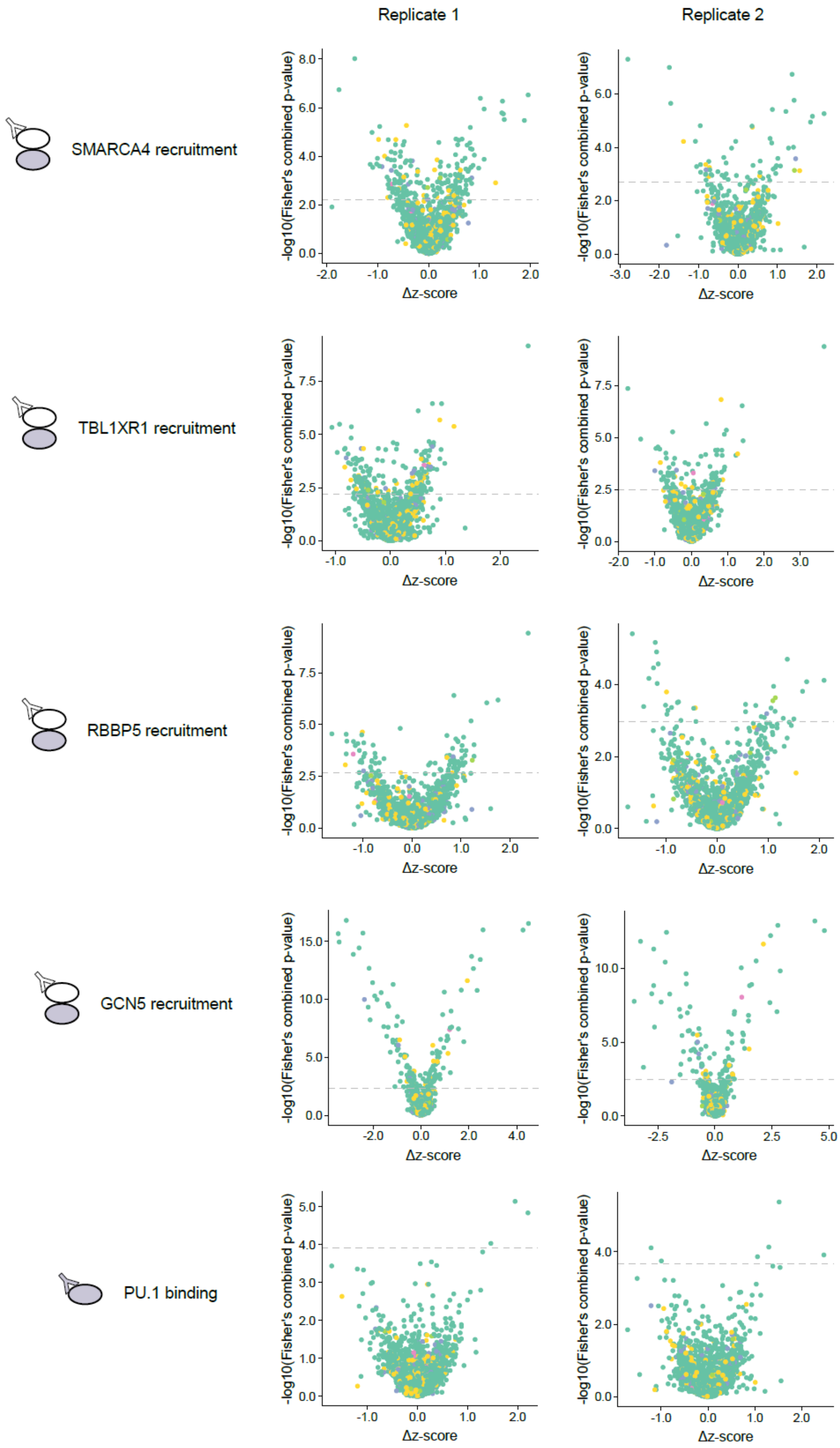
Replicate 1
Replicate 2
SMARCA4 recruitment
TBL1XR1 recruitment
RBBP5 recruitment
GCN5 recruitment
PU.1 binding
-log10(Fisher's combined p-value)
Δz-score

**Supplementary Figure 3. Statistical significance and direction-of-effect for changes in COF recruitment and TF binding across reference and SNP probe pairs screened**. Rows represent volcano plots obtained for different COFs (SMARCA, TBL1XR1, RBBP5, GCN5) and TF PU.1. Left column shows the volcano plots obtained in a first replicate and right column shows the volcano plots obtained in a technical replicate experiment. Statistical significance threshold for each experiment ($q < 0.05$, see Methods) is shown as a grey dashed line. COF: transcriptional cofactor, TF: transcription factor.

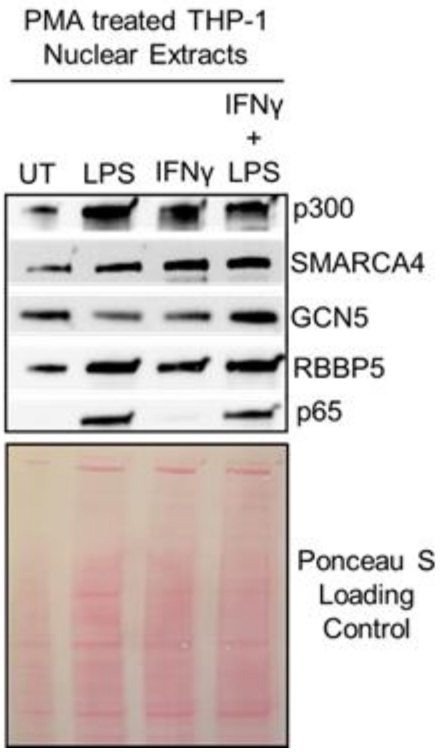


**Supplementary Figure 4. Western blot of PMA treated THP-1 nuclear extracts.** The protein expression levels of p300, SMARCA4, GCN5, RBBP5, and p65 of PMA treated THP-1 cells were evaluated by western blotting. 30ug of nuclear extract were loaded for all samples. PMA treated THP-1 cells were treated with LPS for 45 min to induce p65 expression. PMA treated THP-1 cells were treated with IFNγ for 3 h to prime the immune response. PMA treated THP-1 cells were treated with IFNγ for 1 h and LPS were treated with IFNγ for 2 h followed by LPS stimulation for 45 min. Ponceau S staining was used as a loading control. UT: untreated, LPS: Lipopolysaccharide, IFNγ: Interferon gamma, IFNγ + LPS: Interferon gamma and lipopolysaccharide.
